# Symbiotic bacteria, immune-like sentinel cells, and the response to pathogens in a social amoeba

**DOI:** 10.1101/2023.05.27.542568

**Authors:** Trey J. Scott, Tyler J. Larsen, Debra A. Brock, So Yeon Stacey Uhm, David C. Queller, Joan E. Strassmann

**Affiliations:** Departmet of Biology, Washington University in St. Louis

**Keywords:** sentinel cells, symbiosis, pathogens, *Dictyostelium*, *Paraburkholderia*

## Abstract

Some endosymbionts living within a host must modulate their hosts’ immune systems in order to infect and persist. We studied the effect of a bacterial endosymbiont on a facultatively multicellular social amoeba host. Aggregates of the amoeba *Dictyostelium discoideum* contain a subpopulation of sentinel cells that function akin to the immune systems of more conventional multicellular organisms. Sentinel cells sequester and discard toxins from *D. discoideum* aggregates and may play a central role in defense against pathogens. We measured the number and functionality of sentinel cells in aggregates of *D. discoideum* infected by bacterial endosymbionts in the genus *Paraburkholderia.* Infected *D. discoideum* produced fewer and less functional sentinel cells, suggesting that *Paraburkholderia* may interfere with its host’s immune system. Despite impaired sentinel cells, however, infected *D. discoideum* were less sensitive to ethidium bromide toxicity, suggesting that *Paraburkholderia* may also have a protective effect on its host. By contrast, *D. discoideum* infected by *Paraburkholderia* did not show differences in their sensitivity to two non-symbiotic pathogens. Our results expand previous work on yet another aspect of the complicated relationship between *D. discoideum* and *Paraburkholderia*, which has considerable potential as a model for the study of symbiosis.

## INTRODUCTION

Microbes live in a world replete with other microbes with which they must interact. The most intimate interactions are symbioses, in which unlike organisms live closely associated with or even inside of one another. Symbioses can have many different effects on the participants’ fitness, abilities, and evolutionary fate. Many symbioses enable organisms to survive in ways otherwise beyond them – some symbionts expand the resources their partners can use (Hosokawa et al. 2010; Moran, McCutcheon, and Nakabachi 2008), increase their resistance to abiotic stress (Singh, Gill, and Tuteja 2011),or protect them from hostile organisms (Brownlie and Johnson 2009; Gerardo and Parker 2014). Some of the most dramatic examples of symbiosis have had enormous impacts on the history of life, from enabling the development of complex multicellular organisms to shifting the composition of the planet’s atmosphere on a grand scale (Sagan 1967; Martin, Garg, and Zimorski 2015; Whatley 1993).

However, the line between friend and foe can be blurry. Wherever organisms come together, there will be conflict, even within the most intimate and long-lasting friendships (Budar, Touzet, and De Paepe 2003; Aanen, Spelbrink, and Beekman 2014). Many beneficial symbioses are thought to have evolved from initially antagonistic relationships between partners that later buried the proverbial hatchet (Weiblen and Treiber 2015). Other symbioses may be neither clearly antagonistic nor clearly mutualistic, but rather involve partners that either help or hurt one another depending on the environmental context or the genotypes of the partners involved (Leung and Poulin 2008). Because symbiotic partners do not always have each other’s best interests at heart, it may sometimes be necessary for symbionts to defend themselves from their partners, even in apparently mutualistic symbioses (Gerardo, Hoang, and Stoy 2020; Gross et al. 2009). Conflict will often drive a need for symbionts to modify their own behavior – or that of their partners – to coexist stably, and how symbiotic partners attune to one another is of special interest to understanding how symbioses start and which symbioses endure.

In this study, we focus on the social amoeba *Dictyostelium discoideum* and endosymbiotic bacteria in the genus *Paraburkholderia. D. discoideum* is a normally unicellular eukaryote with a long history of being used as a model organism for scientists interested in its many multicellular behaviors. *D. discoideum* and its relatives are facultatively multicellular organisms that spend most of their time as single, amoeboid cells, moving through forest soil, hunting bacteria, and reproducing vegetatively (Bonner 1944; Kessin 2001; Strassmann and Queller 2011). In adverse conditions, however, *D. discoideum* cells will aggregate into multicellular groups and undergo a sophisticated developmental process to form first a slug-like body that can travel to find a suitable site and then a fruiting body with which to remain dormant until they can disperse to greener pastures (Huss 1989; smith, Queller, and Strassmann 2014). Formation of the fruiting body requires the sacrifice of some cells within the aggregate to form a stalk to hold the other cells aloft. These sacrificial stalk cells are akin to the somatic cells of more conventional multicellular organisms, performing some non-reproductive function (in this case, a structural one) so that other cells can reproduce. Many aspects of *D. discoideum’s* development and evolution have been the focus of studies within a variety of fields (Annesley and Fisher 2009; Bozzaro 2019).

Though they are the most obvious, the cells that die to form *D. discoideum’s* stalk during the last stage of its development are not the only cells that perform somatic functions within *D. discoideum* aggregates. During the slug stage that precedes fruiting, another, smaller subpopulation of ‘sentinel cells’ circulate within the aggregate collecting foreign bacteria and toxins, and are eventually sloughed off the slug and left behind prior to fruiting (Chen, Zhuchenko, and Kuspa 2007). Though the adaptive significance of the sentinel cells’ efforts to clear bacteria and toxins from the slug is not known, it seems likely they serve as a primitive immune system for *D. discoideum*.

*D. discoideum* interacts with a wide variety of soil bacteria in nature (Brock et al. 2018; Haselkorn et al. 2019). Some are its prey, some are its pathogens, and some lie somewhere in between. Among this latter category are *Paraburkholderia agricolaris, P. hayleyella,* and *P. bonniea,* which persistently infect *D. discoideum* cells as intracellular passengers (Brock et al. 2020). Though in many respects *Paraburkholderia* acts as a pathogen, reducing the apparent fitness of its host, infection by *Paraburkholderia* imbues *D. discoideum* with the ability to carry prey bacteria with it throughout the social stages of its life cycle (DiSalvo et al. 2015). This bacterial carriage can enable *D. discoideum* to disperse to prey-impoverished environments not available to uninfected *D. discoideum.* Under the right conditions, therefore, *Paraburkholderia* may have a net benefit for its hosts and behave more like a mutualistic symbiont rather than a pathogen.

The interaction between *D. discoideum* and these *Paraburkholderia* species has both positive and negative consequences for both participants, and is a rising model system in the study of the evolution of interspecific interactions (Scott, Queller, and Strassmann 2022a; Garcia et al. 2019; Brock et al. 2011; Brock et al. 2013; DiSalvo et al. 2015). A heretofore largely unexplored direction, however, is how infection by *Paraburkholderia* may modulate *D. discoideum’s* interactions with other bacteria.

In this study we examine *Paraburkholderia’s* effect on *D. discoideum* sentinel cells, and explore the consequences of these effects on *D. discoideum’s* interactions with other bacterial pathogens that it might encounter in its natural habitat. This work builds on a previous study where we discovered that wild *D. discoideum* isolates infected by *Paraburkholderia* produce fewer sentinel cells than uninfected isolates (Brock, Callison, et al. 2016). If in fact sentinel cells perform an important immune function for *D. discoideum,* it seems intuitive that any disruption of sentinel cell function could render *D. discoideum* hosts more sensitive to toxins and pathogens. However, earlier work suggests that *Paraburkholderia* infection may actually increase hosts’ resistance to ethidium bromide, despite interfering with the sentinel cells that would otherwise remove such toxins from *D. discoideum* aggregates. We hypothesized that it might provide a similar protective effect against intracellular pathogens that might otherwise threaten it and its hosts’ fitness. In this study, we take advantage of recent advances in our understanding of the diversity of *Paraburkholderia* to further explore the consequences of its relationship with its *D. discoideum* hosts.

## MATERIALS AND METHODS

### Culture conditions for *D. discoideum* clones and bacteria symbionts

We used *D. discoideum* clones collected from Virginia, Texas, and North Carolina. We grew *D. discoideum* clones from frozen spore stocks on SM/5 nutrient agar plates (2 g glucose, 2 g Oxoid bactopeptone, 2 g Oxoid yeast extract, 0.2 g MgSO_4_, 1.9 g KH_2_PO_4_, 1 g K_2_HPO_4_ and 15.5 g agar per liter DDH_2_O) and food bacteria *Klebsiella pneumoniae* at room temperature (22°C). We obtained *K. pneumoniae* from the Dicty Stock Center. *K. pneumoniae* was streaked onto SM/5 plates from frozen stocks and allowed to grow until stationary phase. We prepared *K. pneumoniae* bacterial suspensions with an optical density (OD) of A600 1.50 in KK2 buffer (2.25 g KH_2_PO_4_ and 0.67 g K_2_HPO_4_ per liter DDH2O) using a BioPhotometer (Eppendorf, New York). The *D. discoideum* clones and specific symbionts used in these experiments are included in **Table 1**. We removed (cured) *Paraburkholderia* from the infected clones using either ampicillin-streptomycin or tetracycline antibiotic treatment. We verified *Paraburkholderia* removal using PCR with *Paraburkholderia* specific primers (DiSalvo et al. 2015).

**Table 1.**
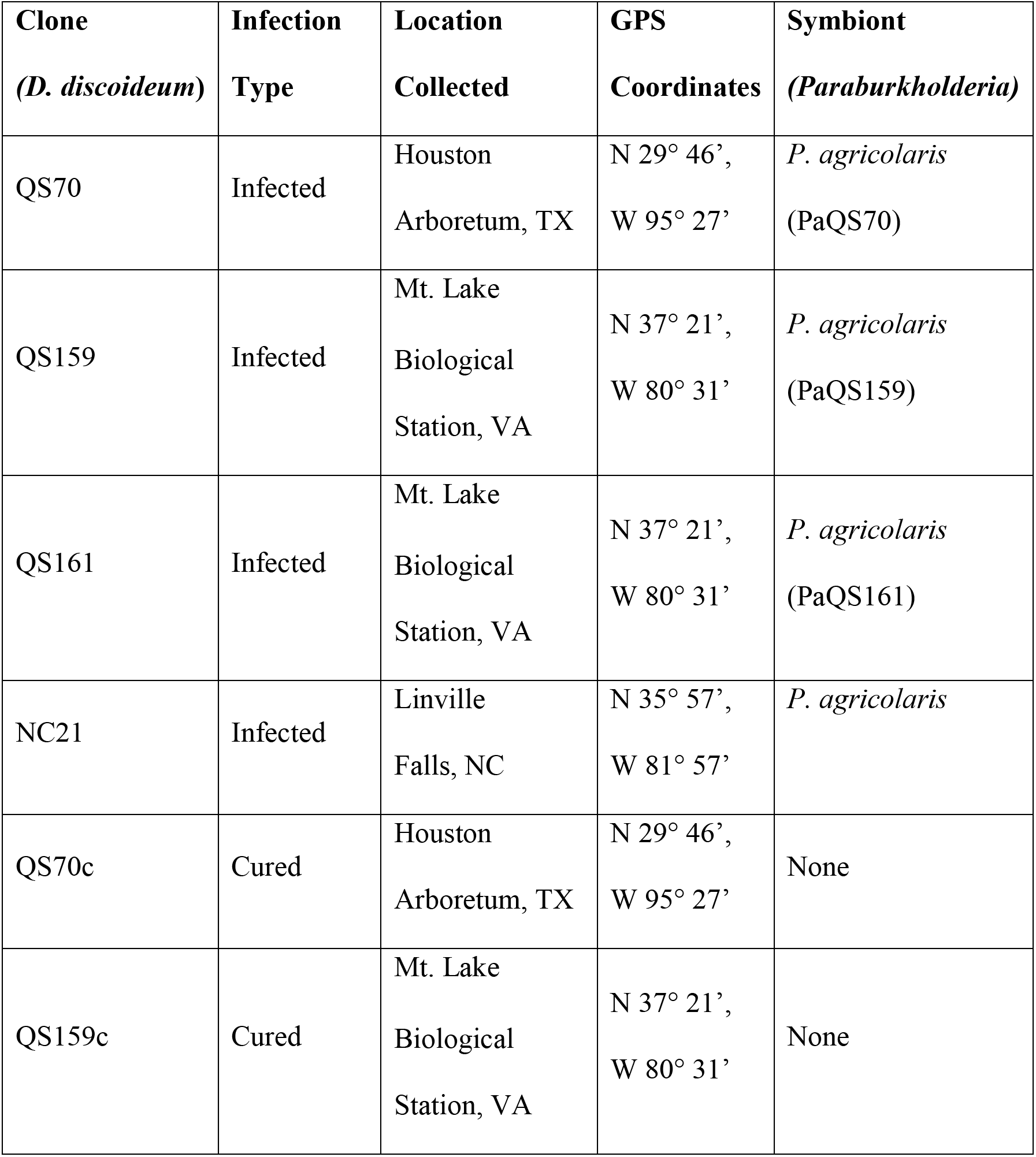
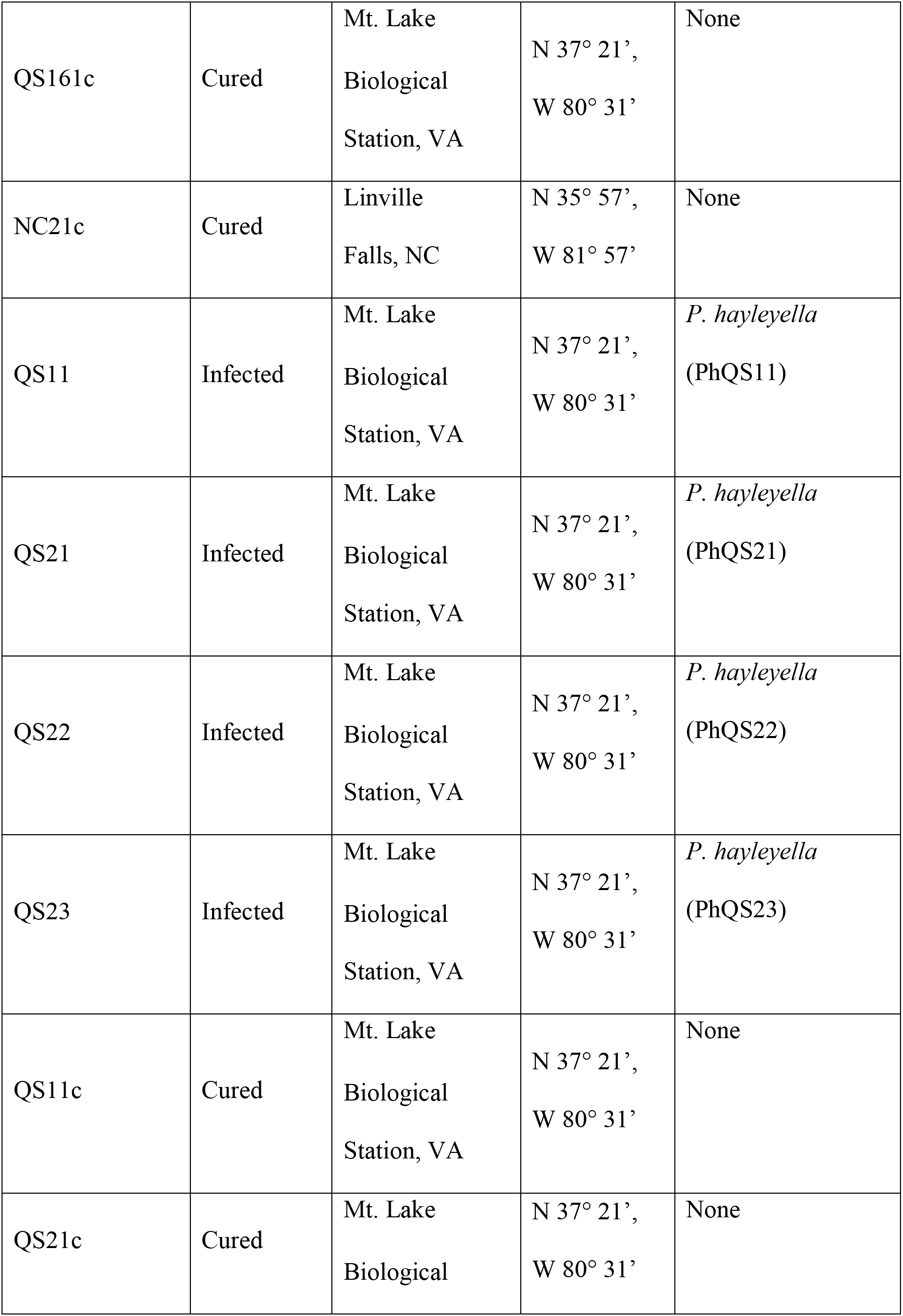
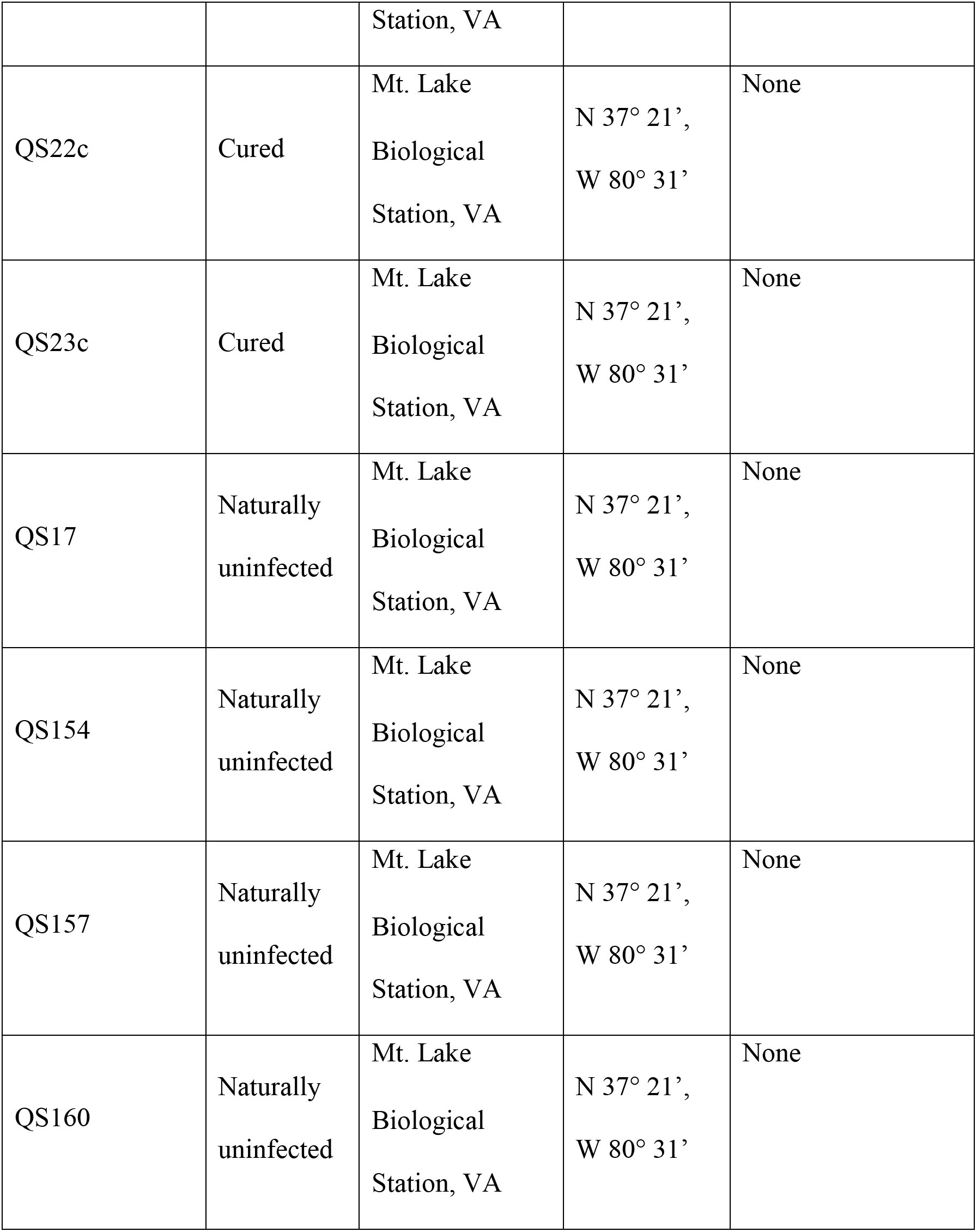
Table of strains.

| **Clone**<br>*(D. discoideum)* | **Infection**<br>**Type** | **Location**<br>**Collected** | **GPS**<br>**Coordinates** | **Symbiont**<br>*(Paraburkholderia)* |
| --- | --- | --- | --- | --- |
| QS70 | Infected | Houston<br>Arboretum, TX | N 29° 46',<br>W 95° 27' | *P. agricolaris*<br>(PaQS70) |
| QS159 | Infected | Mt. Lake<br>Biological<br>Station, VA | N 37° 21',<br>W 80° 31' | *P. agricolaris*<br>(PaQS159) |
| QS161 | Infected | Mt. Lake<br>Biological<br>Station, VA | N 37° 21',<br>W 80° 31' | *P. agricolaris*<br>(PaQS161) |
| NC21 | Infected | Linville<br>Falls, NC | N 35° 57',<br>W 81° 57' | *P. agricolaris* |
| QS70c | Cured | Houston<br>Arboretum, TX | N 29° 46',<br>W 95° 27' | None |
| QS159c | Cured | Mt. Lake<br>Biological<br>Station, VA | N 37° 21',<br>W 80° 31' | None |
| QS161c | Cured | Mt. Lake<br>Biological<br>Station, VA | N 37° 21',<br>W 80° 31' | None |
| NC21c | Cured | Linville<br>Falls, NC | N 35° 57',<br>W 81° 57' | None |
| QS11 | Infected | Mt. Lake<br>Biological<br>Station, VA | N 37° 21',<br>W 80° 31' | *P. hayleyella*<br>(PhQS11) |
| QS21 | Infected | Mt. Lake<br>Biological<br>Station, VA | N 37° 21',<br>W 80° 31' | *P. hayleyella*<br>(PhQS21) |
| QS22 | Infected | Mt. Lake<br>Biological<br>Station, VA | N 37° 21',<br>W 80° 31' | *P. hayleyella*<br>(PhQS22) |
| QS23 | Infected | Mt. Lake<br>Biological<br>Station, VA | N 37° 21',<br>W 80° 31' | *P. hayleyella*<br>(PhQS23) |
| QS11c | Cured | Mt. Lake<br>Biological<br>Station, VA | N 37° 21',<br>W 80° 31' | None |
| QS21c | Cured | Mt. Lake<br>Biological | N 37° 21',<br>W 80° 31' | None |
|  |  | Station, VA |  |  |
| QS22c | Cured | Mt. Lake<br>Biological<br>Station, VA | N 37° 21',<br>W 80° 31' | None |
| QS23c | Cured | Mt. Lake<br>Biological<br>Station, VA | N 37° 21',<br>W 80° 31' | None |
| QS17 | Naturally<br>uninfected | Mt. Lake<br>Biological<br>Station, VA | N 37° 21',<br>W 80° 31' | None |
| QS154 | Naturally<br>uninfected | Mt. Lake<br>Biological<br>Station, VA | N 37° 21',<br>W 80° 31' | None |
| QS157 | Naturally<br>uninfected | Mt. Lake<br>Biological<br>Station, VA | N 37° 21',<br>W 80° 31' | None |
| QS160 | Naturally<br>uninfected | Mt. Lake<br>Biological<br>Station, VA | N 37° 21',<br>W 80° 31' | None |

### Visualizing Sentinel Cells in Slug Trail Assay

To determine if *D. discoideum* sentinel cell numbers are reduced by *Paraburkholderia* presence, we used four clones colonized with *P. agricolaris* and four clones colonized with *P. hayleyella*. These clones include QS70, QS159, QS161, and NC21 for *P. agricolaris* and QS11, QS21, QS22, and QS23 for *P. hayleyella*. We used the same eight clones cured of their *Paraburkholderia* infections as our uninfected control to compare against the infected *D. discoideum* clones.

We adapted methods from (Brock, Callison, et al. 2016) to visualize sentinel cells by staining *D. discoideum* isolates with ethidium bromide (EtBr), an intercalating agent that interferes with nucleic acid synthesis and commonly used as a fluorescent tag (Waring 1965). 50 × 15 mm petri plates were used to prepare non-nutrient agar petri plates (9.9 g KH_2_PO_4_ monobasic, 1.78 g Na_2_HPO_4_ dibasic and 15.5 g agar per liter DDH_2_O) containing 1.0 µg per ml EtBr. Sixty mL of non-nutrient agar was poured first and allowed to set. We laid three microscope slides (3” × 1” × 1 mm) touching each other on top of the cooled agar. Then the slides were embedded in the agar by adding 25 ml of the same non-nutrient agar on top of the slides.

To set up the migration plates, we prepared a concentrated *K. pneumoniae* bacteria suspension. We used an overnight bacterial culture started from a single colony, and grown in Luria Broth (10 g tryptone, 5 g Oxoid yeast extract and 10 g NaCl per liter DDH_2_O) shaking in 25°C. Next day, we pelleted the overnight culture by centrifugation at 10,000*g* for 5 min at 4°C discarding the supernatant. The bacterial pellet was re-suspended and washed in KK2 buffer (2.25 g KH_2_PO_4_ and 0.67 g K_2_HPO_4_ per liter DDH_2_O). We resuspended the final pellet in a small volume of KK2. The bacterial suspension was diluted accordingly to obtain an optical density (OD) A_600_ of 35.00 using a BioPhotometer (Eppendorf, New York).

We collected *D. discoideum* spores in KK2 buffer and determined the spore count using a hemacytometer and light microscope. We suspended 2.0 × 10^5^ spores in 200 µl of prepared bacterial suspension, and 50 µl of the mixture was dispensed in a line parallel to the embedded microscope slides on one edge of the plate. The spore mixture was allowed to dry. The plate was wrapped in aluminum foil with a small hole poked on the opposite side of the line where the spores were deposited. The plates were placed so the hole faced a source of light, towards which the slugs would migrate, and were stored at room temperature (22°C) for 168 hours to allow for sufficient slug migration across the ethidium bromide starving agar petri plate.

To visualize sentinel cells present in slug trails, we excised the embedded microscope slides and placed microscope slide coverslips (24 × 60 mm) on top of the slug trails on the agar. These trails were imaged using a Nikon A1Si laser scanning confocal microscope (Nikon, Tokyo), at 10× magnification under UV light (Texas Red filter, λ = 561.3 nm). We used the “Scan Large Image” function in the software program (NIS Elements Advanced Research version 4.12.01) to capture an image of the slug trails over a large span of area. This function captures multiple images over the selected area and stitches the images together (10% blend). We then used the “Annotations and Measurements” tool to measure the length of the sectioned trails from which sentinel cells were counted. Present in the slug trails were both single and clumped groups of sentinel cells. We counted sentinel cells in clumped groups as an estimate based on the size of one sentinel cell.

### Colonizing Uninfected hosts with *Paraburkholderia* and Migration Assay

To test if colonization with *Paraburkholderia* affects sentinel cell number, we first needed to determine what proportion of *Paraburkholderia* to mix with the food bacteria *K. pneumoniae*. Previously, we had determined that sufficiently high infectious doses of *Paraburkholderia* are toxic enough to prevent hosts from forming slugs or fruiting bodies (data not shown).

We prepared bacterial suspensions of *K. pneumoniae, P. agricolaris* BaQS159, and *P. hayleyella* BhQS11 at OD_600_=35.00 using the method described above. We then combined suspensions of food bacteria and either *P. agricolaris* or *P. hayleyella* at different ratios to compare the effects of different infectious doses. We performed 10-fold serial dilutions to test for slug formation from 1% *Paraburkholderia* + 99% *K. pneumoniae* down to 0.0001% *Burkholderia* + 99.9999% *K. pneumoniae.* The low percentage of *Paraburkholderia* was achieved by diluting the *Paraburkholderia* bacterial suspension down to a lower optical density such that sufficient volume could be pipetted. We did not obtain adequate slug formation until we reduced the percentage of *Paraburkholderia* to 0.001% and 0.0001%. We followed the same steps to plate spores on ethidium bromide plates as described in the visualizing sentinel cells assay above. We used 100% *K. pneumoniae* as our control.

After the slugs were allowed to migrate and fruit, we tested sori to determine if bacteria carriage was induced successfully in naturally uninfected clones. We adapted methods of the spot test described in (Brock et al. 2011). From the fruiting bodies formed at the ends of the slug trails, we randomly picked up individual sori using a filtered pipet tip. Each sorus was transferred onto SM/5 nutrient agar plates as individual spots. We incubated the plates at room temperature (22°C) for two days, examined for bacteria growth, and recorded the number of positive spots of bacterial growth.

### Bead Uptake Assay

To determine if Paraburkholderia infected sentinel cells are able to function as well as those from uninfected hosts, we counted the uptake of 0.5um fluorescent latex beads in sentinel cells present in amoebae from dissociated slugs. We used three types of host clones consisting of four naturally uninfected, three infected by *P. agricolaris* infected, and three infected by *P. hayleyella* infected. We counted the number of beads present in each of ten sentinel cells for each *D. discoideum* clone in each set for a total of 100 sentinel cells.

### Pathogen fitness assay

To assess fitness effects associated with carriage of *Paraburkholderia*, we chose two pathogenic bacteria species known to infect *D. discoideum* intracellularly (*Staphylococcus aureus* Rosenbach (Wichita) ATCC 29213, and *Salmonella enterica* ATCC 14028) (Sillo et al. 2011; Mesquita et al. 2017). For this assay, we tested four hosts naturally infected with *P. agricolaris*, four hosts cured of their *P. agricolaris*, four hosts naturally infected with *P. hayleyella*, four hosts cured of their *P. hayleyella*, and four naturally uninfected hosts. See **Table 1** for specific clone identities. Each clone was grown on either 100% *S. aureus* (gram-positive pathogen), *S. enterica* (gram-negative pathogen), or *K. pneumoniae* (good-food control). We used total spore production as our measure of host fitness. To set up each assay, we plated 2 × 10^5^ spores of each clone in each condition onto SM/5 agar plates in triplicate. All clones formed fruiting bodies within 2-3 days. We collected spores separately from two of the plates five days after fruiting using the method previously described in (Brock et al. 2011). Briefly, we collected spores by washing plates with KK2 buffer supplemented with 0.01% NP-40 alternative (Calbiochem). Then, we counted spores using a hemacytometer and a light microscope.

### Statistical Analyses

We performed statistical analyses in R (version 3.6.3). To compare the means of different groups, we fit models followed by pairwise contrasts calculated with the *emmeans* package (Lenth 2022) using fdr to adjust for multiple comparisons. Sentinel cell data was collected from slug trails with multiple measures from each trail and multiple trails for each clone. To account for this nested structure of sentinel cell counts, we included the random effects of trail nested within clone in linear mixed models (LMM) using the nlme package (Pinheiro and Bates 2006). We also log transformed sentinel cell counts to reduce the skew from high counts. For our bead results, we fit generalized linear mixed models (GLMM) in the lme4 package (Bates et al. 2015) with a Poisson link function and clone as a random effect.

To measure how pathogens affected spore production, we used a generalized least squares (GLS) model. Because infections with different pathogens resulted in groups with different variances, we weighted observations using the varIdent function in the nlme package. To re-analyze the spore data in Brock et al. 2016, we used a LMM. To account for the two technical replicates used in this experiment, we included clone as a random effect. We scaled spore production values by subtracting the mean and dividing by the standard deviation.

For both our pathogen and ethidium bromide models, we estimate effects relative to the control condition (grown with *K. pneumoniae* food bacteria or without ethidium bromide), where hosts are not expected to suffer reduced spore production. We are thus measuring the effect of some stressor (infection or toxin) relative to healthy hosts. We are mostly interested in the interaction effects between *Paraburkholderia* infection status (infected or cured) and stressors (pathogen infection or toxins). These effects would indicate that Paraburkholderia infection (or curing) protects or causes increased harm for hosts when exposed to a stressor.

## RESULTS

### Are symbiont *Paraburkholderia* the causal agents for immune-like sentinel cell number reduction?

*Paraburkholderia* infection is known to cause numerous phenotypes in *D. discoideum* hosts, including conferring the ability to carry food bacteria (DiSalvo et al. 2015; Brock, Jones, et al. 2016) and a reduction in predation ability (Scott, Queller, and Strassmann 2022b). A prior study showed that *D. discoideum* clones naturally infected by *Paraburkholderia* had fewer sentinel cells than naturally uninfected *D. discoideum* clones (Brock, Callison, et al. 2016). However, it is not yet clear whether *Paraburkholderia* infection specifically causes the reduction in sentinel cell number. We tested whether *Paraburkholderia* infection was responsible for changing the number of sentinel cells by experimentally curing infected hosts and infecting naturally uninfected hosts.

Both curing hosts of their *Paraburkholderia* and infecting naïve hosts showed that *Paraburkholderia* are responsible for changes to sentinel cells. Curing hosts of *P. agricolaris* infections increased the number of sentinel cells by 35 percent (ratio = 0.649, se = 0.072, df = 113, p = 0.002; **Figure 1 A-B**). Curing hosts of *P. hayleyella* infections increased the number of sentinel cells by 40 percent (ratio = 0.600, se = 0.0664, df = 113, p < 0.001). Infecting naturally uninfected (naïve) hosts reduced the number of sentinel cells by 50 percent or more for both 0.001% and 0.0001% infection doses (**Figure 1C**). Infecting hosts with *P. agricolaris* at 0.0001% (ratio = 0.389, se = 0.042, df = 129, p < 0.001) and 0.001% (ratio = 0.398, se = 0.043, df = 129, p < 0.001) resulted in about 60 percent fewer sentinel cells. Infecting hosts with *P. hayleyella* at 0.0001% (ratio = 0.504, se = 0.055, df = 129, p < 0.001) and 0.001% (ratio = 0.468, se = 0.051, df = 129, p < 0.001) resulted in about 50 percent fewer sentinel cells. Infecting hosts with different doses did not affect sentinel cell number for hosts infected with *P. agricolaris* (ratio = 0.979, se = 0.105, df = 129, p = 0.845) or hosts infected with *P. hayleyella* (ratio = 1.077, se = 0.117, df = 129, p = 0.550).

**Figure 1.**
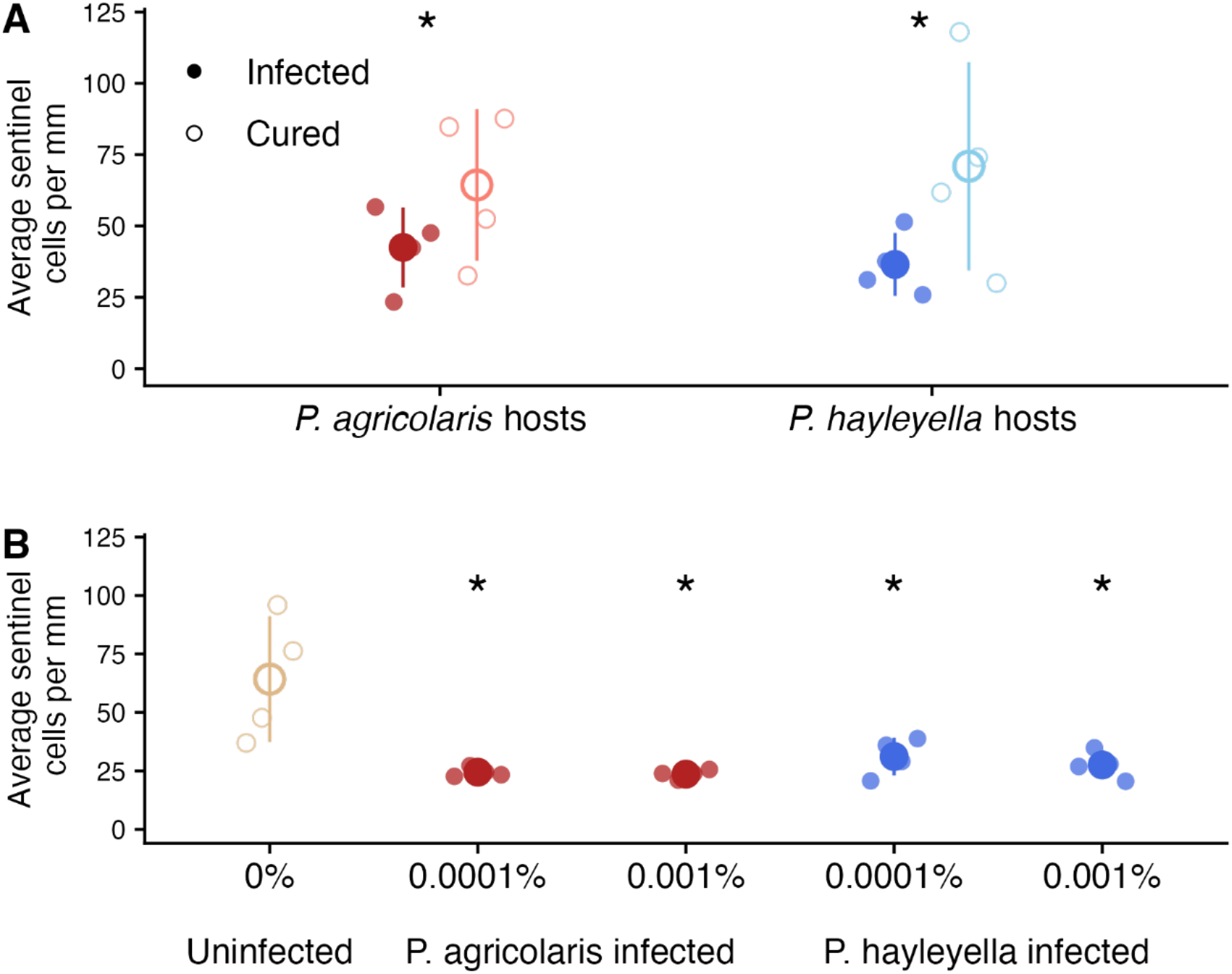
*Paraburkholderia* infection lowers the number of sentinel cells in *D. discoideum* hosts. **A)** Infected *D. discoideum* clones have significantly fewer sentinel cells than the same clones cured of *Paraburkholderia* by antibiotics. The small points represent the average number of sentinel cells counted per mm in individual trails for each clone. **B)** Naturally uninfected *D. discoideum* that are infected with *Paraburkholderia* have fewer sentinel cells even when infected at extremely low doses. Large points and lines show the mean and standard deviation. Asterisks show significant differences (p ≤ 0.05). Statistical comparisons in B are relative to uninfected controls.

### Is the function of sentinel cells from infected hosts impaired?

Because hosts infected with *Paraburkholderia* have fewer sentinel cells (**Figure 1**), we suspected *Paraburkholderia* infection may also impact sentinel cell function. To test sentinel cell function, we measured the number of beads that sentinel cells were able to phagocytose. We found that sentinel cells from *P. agricolaris* infected hosts take up fewer beads (about 17% less) compared to uninfected hosts (**Figure 2**), but this difference was not significant (ratio = 0.830, se = 0.098, p = 0.115) Sentinel cells from *P. hayleyella* infected hosts take up about 30% fewer beads than uninfected hosts (ratio = 0.696, se = 0.085, p = 0.006; Figure 2). Hosts infected by *P. hayleyella* have reduced sentinel cell function.

**Figure 2.**
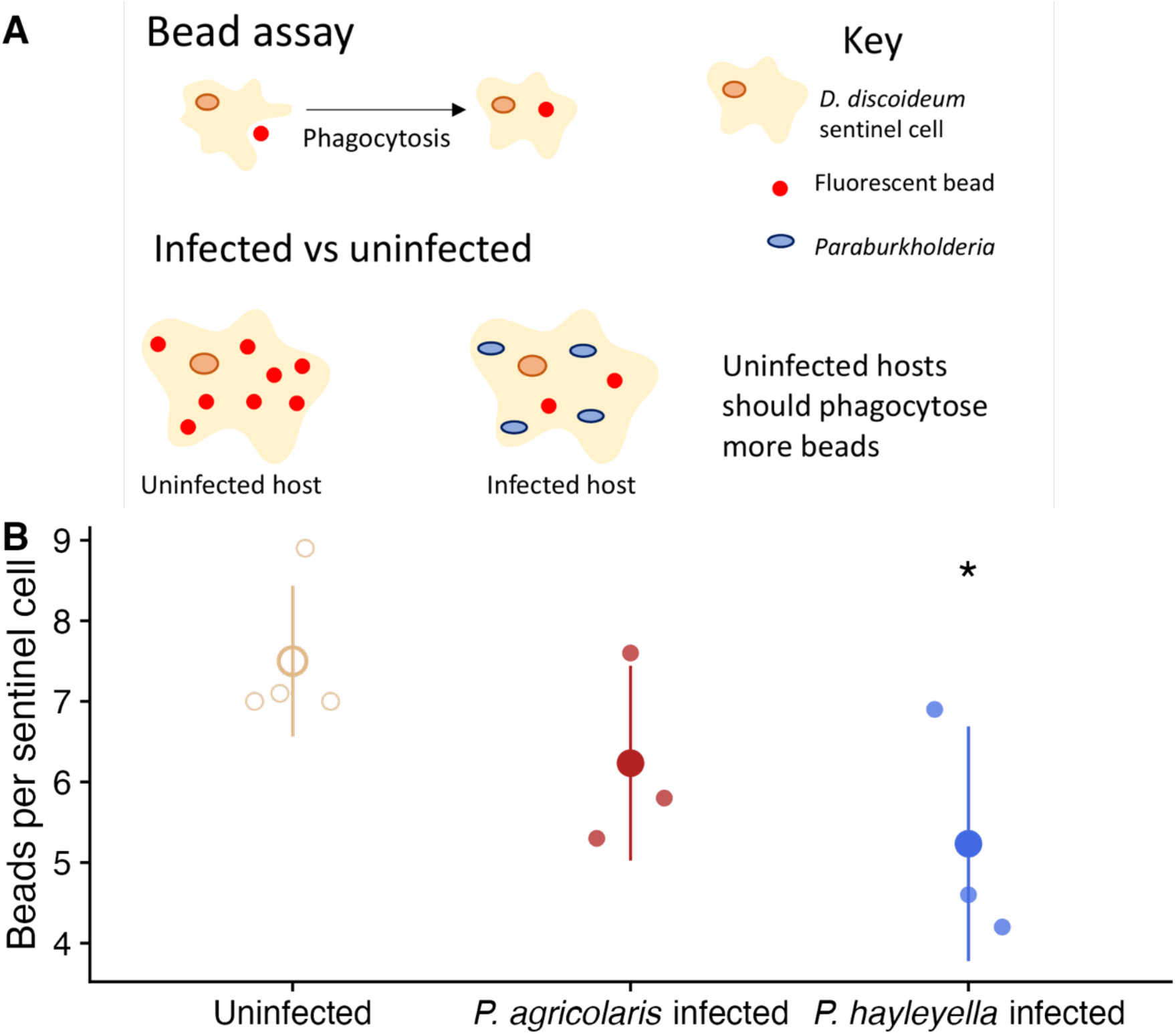
Sentinel cells from infected hosts are less functional than sentinel cells from uninfected hosts. **A)** Schematic of fluorescent bead assay. **B)** Number of beads phagocytosed by uninfected amoebae, amoebae infected with *P. agricolaris*, and amoebae infected with *P. hayleyella*. Small points represent the average number of beads for each clone (counted from ten individual sentinel cells). Large points and lines show the mean and standard deviation. Asterisks show significant differences (p ≤ 0.05) relative to uninfected controls.

### How does Paraburkholderia infection affect host response to toxicity and pathogens?

A prior study found that *Paraburkholderia* infection protected hosts from ethidium bromide toxicity (Brock, Callison, et al. 2016). However, hosts in their natural soil environment are unlikely to come in contact with ethidium bromide. We instead sought a more natural stressor to understand how reduced sentinel cell function affected hosts. We suspected that reduced function of sentinel cells due to *Paraburkholderia* infection may make infected hosts more susceptible to harm from other pathogens.

We first re-analyzed the data from (Brock, Callison, et al. 2016) where infected, uninfected, and cured hosts were grown on starving agar plates with or without toxic ethidium bromide (**Figure 3A**). While the original study included hosts infected by both *P. agricolaris* and *P. hayleyella,* it did not draw a distinction between the two species. (Additionally, the data set only included cured controls for strains infected by *P. hayleyella*). We estimated the effects on host spore production of ethidium bromide, infection, curing, and the interactions between these categories relative to uninfected controls that were not exposed to ethidium bromide. We describe the estimated effects from top to bottom in Figure 3B. We found that ethidium bromide (EtBr) lowered host spore production (Figure 3A&B; estimate = -1.316, 95% CI = [-1.824, - 0.809], df = 59). The harm of ethidium bromide was a similar magnitude as the characteristic cost of infection for both *P. agricolaris* (Pa, estimate = -0.939, 95% CI = [-1.594, -0.284], df = 59) and *P. hayleyella* (Ph, estimate = -1.326, 95% CI = [-1.980, -0.671], df = 59) (DiSalvo et al. 2015). Interestingly, curing hosts of their *P. hayleyella* increased spore production relative to uninfected hosts (Ph cured, estimate = 1.053, 95% CI = [0.458, 1.647], df = 59). Validating the results of (Brock, Callison, et al. 2016) that found a protective effect of *Paraburkholderia* infection, we found that hosts infected with *P. agricolaris* (Pa*EtBr, estimate =1.5547826, 95% CI = [0.629, 2.481], df = 59) or *P. hayleyella* (Ph*EtBr, estimate =1.618, 95% CI = [0.692, 2.544], df = 59) produced more spores when exposed to ethidium bromide than expected from the separate effects of Paraburkholderia infection and ethidium bromide. This effect is due to *Paraburkholderia* infection, at least for *P. hayleyella*, as these hosts cured of their symbionts did not deviate from the additive effects of curing and ethidium bromide (Ph cured*EtBr, estimate = -0.681, 95% CI = [-1.523, 0.160], df = 59).

**Figure 3.**
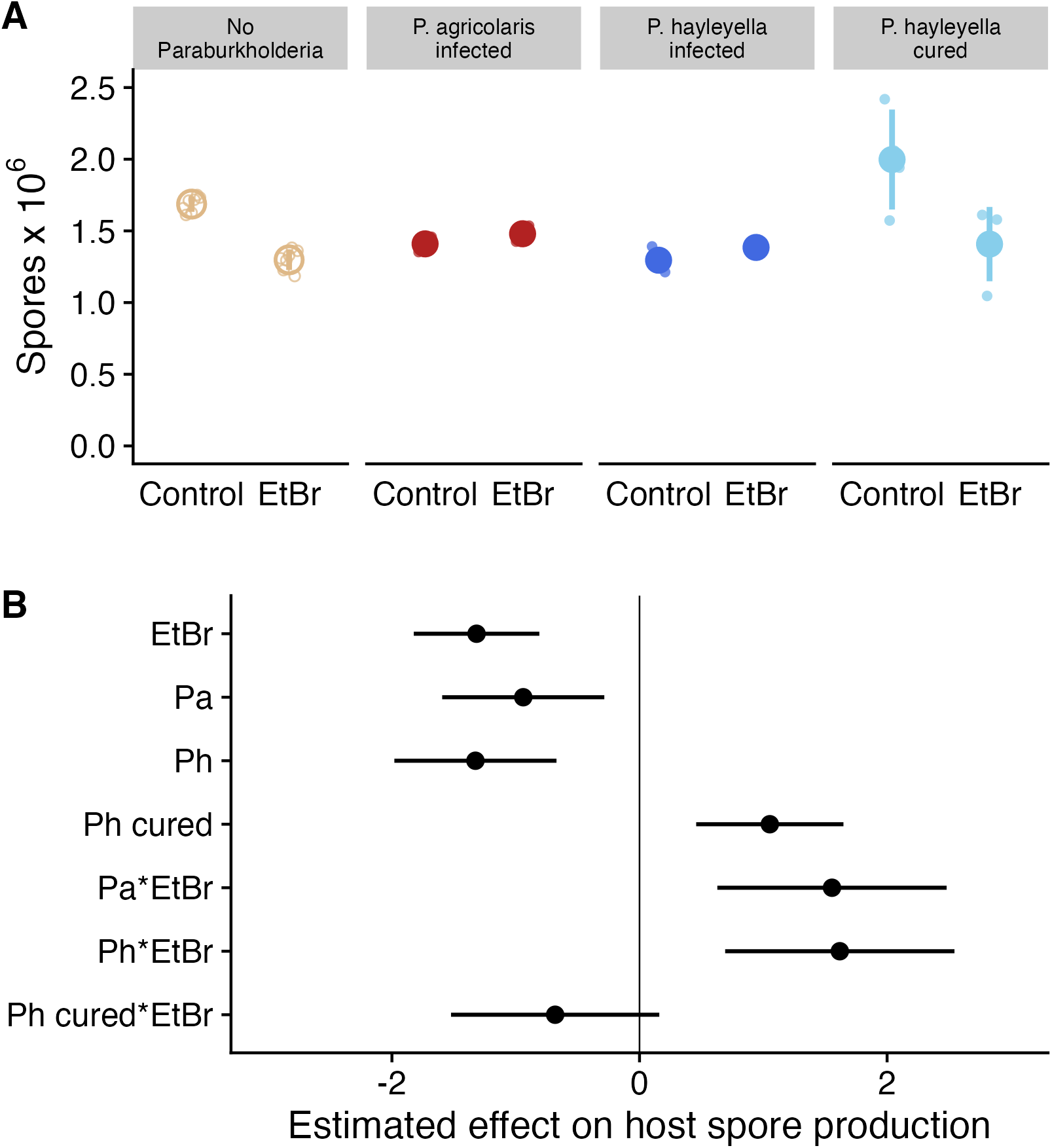
*Paraburkholderia* infection protects hosts from ethidium bromide. **A)** Spore production from hosts with different infection statuses. These data are from Brock et al. 2016 broken up by the species of infecting *Paraburkholderia*. Control *D. discoideum* were grown on starving agar without ethidium bromide. **B)** Estimated effects of ethidium bromide, *Paraburkholderia* infection status, and interactions on host spore production. Host spore production in the model was scaled by subtracting the mean and dividing by the standard deviation.

To address the possibility that *Paraburkholderia* infection makes hosts are more prone to harm from a more natural stressor, we tested spore production in the presence of two pathogenic bacteria, *Salmonella enterica* and *Staphylococcus aureus*. We used *Klebsiella pneumoniae* as our control food bacteria for comparison. To measure the harm of infection with these other pathogens, we counted the spores produced by *D. discoideum* (Figure 4A). We asked two questions: (1) Does infection with pathogens lower *D. discoideum* spore production relative to growing on food bacteria? (2) Does infection with *Paraburkholderia* affect harm to hosts when infected with other pathogens?

**Figure 4.**
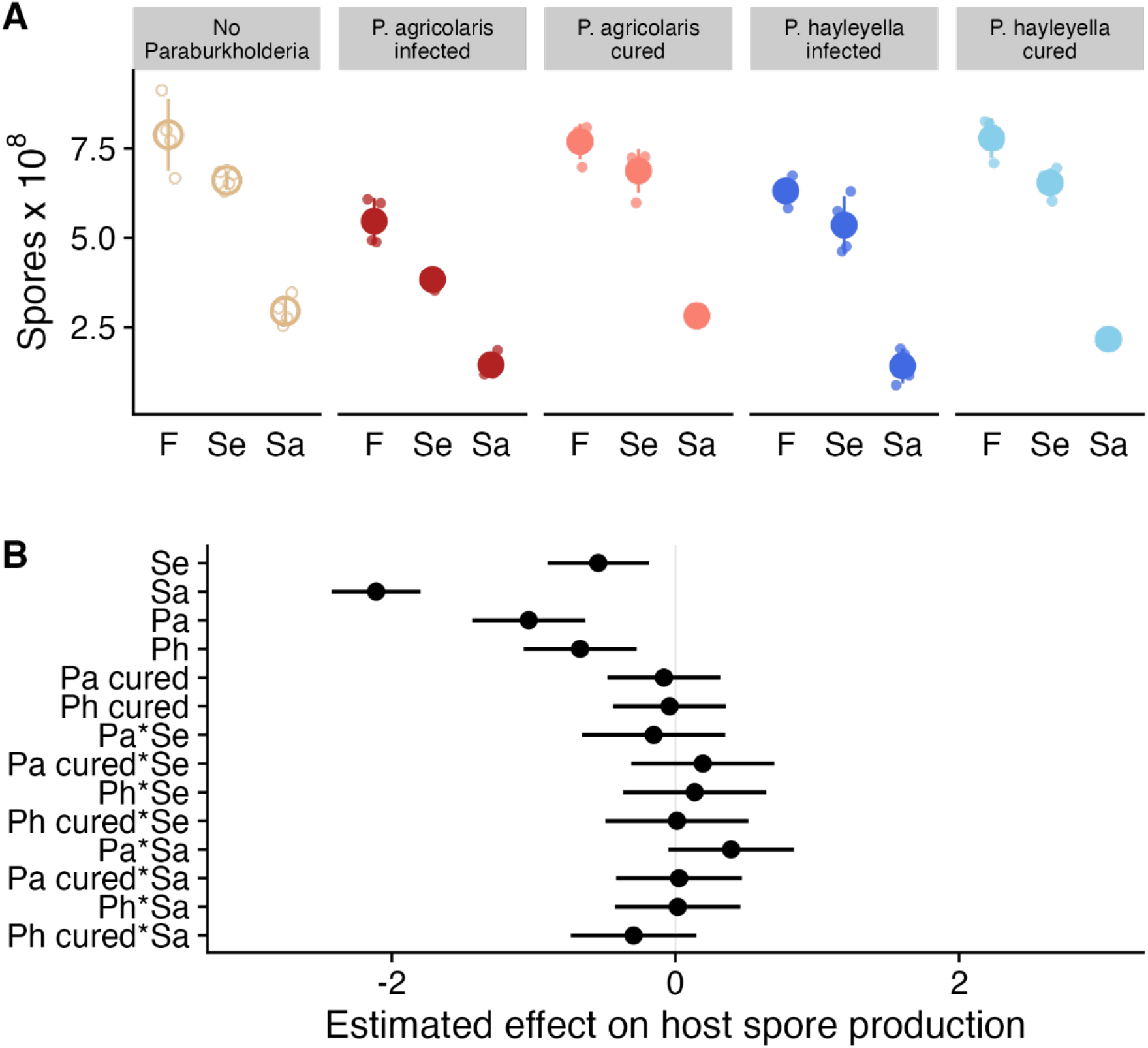
*Paraburkholderia* infection does not protect hosts from pathogenic Salmonella enterica (Se) or Staphylococcus aureus (Sa) bacteria. A) Spore production of hosts when grown on food bacteria (F) or pathogenic *Salmonella enterica* (Se) or pathogenic *Staphylococcus aureus* (Sa).**B**) Estimated effects (points) and 95% confidence intervals of *Paraburkholderia* infection (Pa = *P. agricolaris*; Ph = *P. hayleyella*), pathogen infection (Sa or Se), and coinfection (shown joined by *) on *D. discoideum* spore production. Spore production values were scaled by subtracting the mean and dividing by the standard deviation.

To answer these questions, we fit a generalized least squares model with interaction terms between each pathogen infection status (either *S. enterica* or *S. aureus*) and each *Paraburkholderia* infection status (either *P. agricolaris*, cured of *P. agricolaris*, *P. hayleyella*, or cured of *P. hayleyella*) to *D. discoideum* spore production measures (**Figure 4A**). We used this model to estimate the effects of pathogen infection, *Paraburkholderia* status, and coinfections (interactions) relative to uninfected *D. discoideum* that were grown on only food bacteria (**Figure 4B**). Working down from the top of Figure 4B, we found that *S. enterica* is moderately pathogenic (Se, estimate = -0.544, 95% CI = [-0.901, -0.188], df = 60) while *S. aureus* is highly pathogenic, reducing host spore production almost fourfold (Sa, estimate = -2.110, 95% CI = [-2.423, -1.797], df = 60) relative to *S. enterica*. *Paraburkholderia* infection by *P. agricolaris* (Pa, estimate= -1.034, 95% CI = [-1.432, -0.636], df = 60) or *P. hayleyella* (Ph, estimate =-0.672, 95% CI = [-1.070, -0.273], df = 60) is similarly pathogenic as *S. enterica*, and curing obviated this cost (Figure 4B; cured Pa and cured Ph effects are essentially 0).

Coinfection by *Paraburkholderia* and a pathogen did not increase or decrease harm from pathogens beyond the additive effects of both kinds of infection (interaction terms with *’s overlap 0). These results show that *Paraburkholderia* and pathogen infections both reduce *D. discoideum* spore production, but that *Paraburkholderia* does not make hosts more susceptible to harm by pathogens. On the other hand, *Paraburkholderia* also offers no protection against the pathogens we tested in contrast to the results with ethidium bromide.

## DISCUSSION

In this study, we investigated the effects of *Paraburkholderia* infection on the sentinel cells of *D. discoideum* hosts. Due to sentinel cells’ presumed function as a primitive immune system within *D. discoideum* aggregates, we also explored the consequences of *Paraburkholderia* infection on *D. discoideum*’s sensitivity to toxins and pathogens.

*D. discoideum* aggregates produce a subpopulation of sentinel cells which seem to serve as an innate immune system for the multicellular stages of its life cycle (Chen, Zhuchenko, and Kuspa 2007). Like the cellular immune systems of more familiar multicellular organisms, sentinel cells circulate within *D. discoideum* aggregates, sequestering and disposing of potentially hazardous foreign material like toxins or pathogens. In a previous study, we observed that some wild *D. discoideum* isolates carrying certain *Paraburkholderia* bacteria through their social cycles also produced significantly fewer sentinel cells (Brock, Callison, et al. 2016).

To explore the role of *D. discoideum’s* intracellular symbiont *Paraburkholderia* on its host’s sentinel cells, we compared the number of sentinel cells produced in slugs made by wild *D. discoideum* strains known to harbor *P. agricolaris* or *P. hayleyella* symbionts with slugs made by the same strains after antibiotic treatment. We found that antibiotically treated *D. discoideum*, cured of their normal symbionts, produced significantly more sentinel cells than infected cells. When we then reinfected these cured strains with *P. agricolaris* or *P. hayleyella* in the laboratory, the effect was reversed and fewer sentinel cells were produced. These results establish a causal link between infection by *Paraburkholderia* and the previously observed reduced sentinel cell numbers in infected *D. discoideum* strains. *Paraburkholderia* infections reduced *D. discoideum* sentinel cell production even at very small infectious doses.

In addition to quantifying the reduction in sentinel cell numbers in infected *D. discoideum*, we also tested sentinel cell functionality by measuring sentinel cells’ ability to sequester fluorescently labeled beads. We found that individual sentinel cells produced by infected *D. discoideum* infected sequestered significantly fewer beads than sentinel cells from uninfected *D. discoideum*.

Our results indicate that *D. discoideum* infected by *Paraburkholderia* not only produces fewer sentinel cells, but those that it does produce are less functional. Insofar as sentinel cells serve an important role in clearing potentially damaging foreign substances from multicellular *D. discoideum* aggregates prior to the production of fruiting bodies, we expected infected *D. discoideum* to have impaired immune function and be more sensitive to toxins or pathogens. To test this, we first reanalyzed data measuring the toxic effect of ethidium bromide on spore production in infected and uninfected *D. discoideum* (Brock, Callison, et al. 2016). Uninfected *D. discoideum* exposed to ethidium bromide produces fewer spores during its fruiting stage, presumably reflecting death or reduced functionality of cells within the aggregate. In contrast, however, *D. discoideum* infected by *Paraburkholderia* produced similar numbers of spores with and without the presence of ethidium bromide. While infected *D. discoideum* produce fewer spores overall due to the effects of *Paraburkholderia* infection itself (DiSalvo et al. 2015), they no longer appear to be sensitive to ethidium bromide’s toxic effects.

Sentinel cells may represent an adaptation against threats posed by pathogens – either through infection or via the production of toxins. *D. discoideum* is known to be susceptible to a wide variety of pathogens (Steinert 2011), which, combined with its similarities to human macrophages, has made it a common model for the study of pathogenesis and immunity (Steinert and Heuner 2005). Previous work suggests that sentinel cells likely serve a protective function against pathogens, presumably by sequestering and removing them from *D. discoideum* slugs as they do with ethidium bromide (Chen, Zhuchenko, and Kuspa 2007). Sentinel cells were shown to upregulate their expression of the *tirA* and *tirB* genes, apparent homologs of genes known to be involved in animal and plant innate immunity signaling. Mutant strains lacking functional *TirA* showed increased sensitivity to the virulent pathogen *Legionella pneumophila*.

In light of sentinel cells’ potential immune function, we explored whether *Paraburkholderia’s* effects on sentinel cell number and functionality increased *D. discoideum’s* sensitivity to other pathogens, as would be expected if sentinel cell activity played a key role in immunity. We exposed uninfected amoebae and amoebae carrying *Paraburkholderia* symbionts to *Staphylococcus aureus* and *Salmonella enterica*, two pathogens known to infect *D. discoideum.* Unsurprisingly, *D. discoideum* exposed to either pathogen produced fewer spores, reflecting *S. aureus* and *Sa. enterica’s* virulent effects. Despite its effects on sentinel cell number and functionality, however, infection by *Paraburkholderia* did not have a significant effect on *D. discoideum’s* sensitivity to either tested pathogen.

Our results suggest that *D. discoideum* infected by *Paraburkholderia* experiences reduced sentinel cell production and function. Building upon previous work, this study demonstrates that *Paraburkholderia* itself is responsible for changes in sentinel cells (rather than some strains of *D. discoideum* evolving reduced sentinel cells to facilitate symbiosis or for some other purpose). Naïve *D. discoideum* strains with no known association with *Paraburkholderia* produce fewer sentinel cells when infected, and infected *D. discoideum* sentinel cell production is restored when *Paraburkholderia* symbionts are cleared by antibiotic treatment.

It is unclear whether *Paraburkholderia*’s effect on its host’s sentinel cells is adaptive. Given that some evidence suggests sentinel cells may be involved in clearing bacteria from *D. discoideum* slugs, it is intuitive to speculate that *Paraburkholderia* could benefit from inhibiting them. Alternately, *Paraburkholderia*’s effects on sentinel cells may be pleiotropic consequences of other traits. *Paraburkholderia*’s intracellular lifestyle is likely only possible because it can waylay the mechanisms by which *D. discoideum* phagocytoses and destroys its prey. If phagocytosis is also key to sentinel cell function during the slug stage, whatever mechanism *Paraburkholderia* uses to survive phagocytosis during its host’s vegetative stage may inhibit sentinel cell function as a side effect.

Despite its effect on sentinel cell number and function, *Paraburkholderia* infection also has the apparently paradoxical effect of *reducing* its host’s sensitivity to the toxin ethidium bromide. One possible explanation is that *Paraburkholderia* may neutralize ethidium bromide in a way which compensates for the presumed loss of *D. discoideum*’s own defenses resulting from fewer, less functional sentinel cells. Many bacteria have metabolic capabilities unavailable to eukaryotes, and these form the basis for various other symbioses (Deschamps et al. 2008; Douglas 1998; Belkin, Nelson, and Jannasch 1986). Previous work has suggested that *Paraburkholderia* and *D. discoideum* have a complex relationship with both antagonistic and cooperative elements, and it is likely that whether *Paraburkholderia* is a beneficial partner or a parasite depends on the specific context. If *Paraburkholderia* can protect its host from toxins, then the likelihood of encountering these toxins could play a role in whether *Paraburkholderia* behaves as a cooperative symbiont or a parasite.

In contrast, we did not observe any effect of *Paraburkholderia* infection on *D. discoideum’s* sensitivity to the pathogens *S. aureus* or *Sa. enterica.* A previous study demonstrated that *D. discoideum* mutants lacking a functional *tirA* gene – normally highly expressed in sentinel cells – were extremely sensitive to virulent *L. pneumophila.* This has been taken to imply an important role for sentinel cells in protecting *D. discoideum* from pathogens, but our results cast some doubt on this interpretation. *Paraburkholderia-*driven reductions in sentinel cell number and functionality did not seem to render *D. discoideum* any less capable of surviving *S. aureus* or *Sa. enterica* as would be expected if they served a vital immune function against these pathogens. It may be that sentinel cells’ potential immune function is context specific, or that *D. discoideum* combats different pathogens in different ways. Alternately, perhaps infected *D. discoideum* with fewer and less functional sentinel cells *are* more sensitive to pathogens but are compensated for somehow by the presence of *Paraburkholderia*.

The specific role *D. discoideum’s* sentinel cells in defending against specific pathogens is not yet fully clear, but we consider it likely that they function as a simple immune system for the multicellular portions of *D. discoideum’s* life cycle. Previous studies demonstrate that sentinel cells sequester and discard toxic ethidium bromide, which may suggest a more general role in combatting toxins produced by other microbes in *D. discoideum’s* environment. If sentinel cells do have an immune function, then *Paraburkholderia’s* ability to reduce sentinel cell production and functionality may be an adaptation to actively interfere with its host’s defenses. Regardless of whether *Paraburkholderia* acts as a pathogen or a mutualist, this sort of immune interference has precedence in other systems. Pathogens, perpetually locked in antagonistic coevolutionary arms races with their hosts, have evolved myriad ways of interfering with every part of their hosts’ immune systems (Brodsky and Medzhitov 2009; Finlay and McFadden 2006). While this sort of antagonism might be expected between foes, it is also known to occur in apparently cooperative interactions. Research on other host/symbiont systems suggests that modulation of the host’s immune system – whether imposed by the host itself or by interference from the symbiont – is often critical for establishment and persistence of symbioses (Zug and Hammerstein 2015; Gourion et al. 2015; Cerf-Bensussan and Gaboriau-Routhiau 2010; Jacobovitz et al. 2021; Detournay et al. 2012).

*Paraburkholderia*’s effect on *D. discoideum*’s sentinel cells adds an intriguing element to our understanding of their already complex relationship. Further exploration into the mechanisms by which *Paraburkholderia* infects and persists within *D. discoideum* will help further develop the system as a model for symbiosis, cooperation, and antagonism between microbes.

## Acknowledgements

We thank members of the Queller Strassmann lab for useful advice in the experiments and the writing. Part of this work was a senior thesis by Stacey Uhm. This material is based upon work supported by the National Science Foundation under grant numbers IOS 1656756, DEB 1753743, and DEB 2237266.

## Data, code, and materials

All data and code for this project are publicly available at https://gitlab.com/treyjscott/sentinel_cell_project.

